# Quantifying punctate morphology in mesenchymal stem cells from single cell images provides a definitive quality assessment pipeline for direct correlation to osteogenic lineage differentiation

**DOI:** 10.64898/2026.10.22.760947

**Authors:** Samikshya Purushotham, Triptish Bhattacherjee, Ayona Gupta, Samantha Shrihari, Lakshmi Balasubramanian, Urmi Bandyopadhyay, Uttara Chakraborty

**Affiliations:** Manipal Institute of Regenerative Medicine, Manipal Academy of Higher Education, Manipal, India; Centre for Cellular and Molecular Platforms (C-CAMP), UAS-GKVK Campus, Bellary Road, Bangalore, Karnataka, India

## Abstract

Mesenchymal stem cells (MSCs) exhibit unique immunological properties and are classified as multipotent stem cells. Considering their immunosuppressive and immunoenhancing properties, they have been used adequately for tissue regeneration across allogeneic barriers. MSCs are propagated mainly using in vitro methods of cell culture and thereby exhibit heterogeneity in a wide spectrum. Such stem cell heterogeneity can arise due to donor, tissue and cellular variability. Studies have shown that functional variances in MSCs derived from the same tissue source across different donors may be influenced by age, gender, and physiological condition. In contrast to their mature counterparts, mesenchymal stem cells exhibit a relative resilience to cellular senescence; yet they remain vulnerable to both intrinsic and extrinsic stress. Cellular senescence can be induced by exposure to cellular stress or a reduction in replicative capacity. Cellular structure, telomere length, and senescence-associated β-galactosidase (SA-β-gal) activity are some of the hallmark observations signifying the onset of senescence in cells. Such a biological state in mesenchymal stem cells diminishes the overall effectiveness of the cells as candidates for biotherapeutics. Although MSCs towards their later passages are known to show drop in proliferative properties and in their regenerative potential, direct correlation of passage related onset of cellular senescence and lineage differentiation at the single cell precision remains largely unexplored. To close this gap in stem cell research, we have developed a single cell image-based pipeline exclusively for human MSCs, which addresses immune-heterogeneity features using both the bright field and fluorescence-based images of stem cells. The MSCProfiler or the single cell image-based pipeline can analyze and quantify feature descriptors of the MSCs, across tissues and passages *in vitro*. We have extrapolated the pipeline to address a critical biological state like senescence in MSCs to quantify this multi-variable property of stem cells and check the correlation with cellular differentiation, if any. To address the properties of replicative senescence and study it at the single cell level, we wanted to study the senescence-associated β-galactosidase activity in human dental pulp derived MSCs (hDPSCs). Further to these analyses, we wanted to assess if quantifying the punctate morphology in single stem cells can establish any direct correlation of senescence with osteogenic lineage differentiation in case of the hDPSCs. To achieve our hypothesis, we used the MSCProfiler and extended the pipeline to incorporate the expression profiles of senescence-associated β-galactosidase across the early, mid and late passages of hDPSCs. The pipeline was modified to incorporate bright field image features such as cell size, shape and eccentricity and fluorescence images from the SA-β-galactosidase-substrate reactions. We classified the puncta morphology across passages of hDPSCs and checked for osteogenic lineage differentiation abilities in these MSCs. This could prove to be definitive quality assessment pipeline to assess the biological state of all types of MSCs before their use in allogenic transplantations.

## 1. Introduction

An inevitable physiological consequence of organisms is aging. Diverse acquired and inherited variables drive the intricate cellular and organismal mechanisms that control mammalian aging [1]. Approximately two thirds of the 150,000 individuals who die every day worldwide are due to age-related reasons [2]. The process of aging is among the most significant recognized risk factors for numerous human illnesses [3]. Cellular senescence is a principal factor in the gradual aging phenomenon. Given their remarkable capacity to address many ailments, “stem cell research” is recognized as one of the most promising domains in the treatment of debilitating conditions and tissue regeneration today. From 1998, since Dr. James Alexander Thomson achieved the significant milestone of generating many embryonic stem cell lines from human blastocysts, stem cell research has become a focal point in biomedical research [4].However, there is now strong evidence that stem cells are negatively impacted by the aging process of the body, and that these impacts may be crucial to the pathophysiology of the several diseases linked to aging [5] such as cardiovascular disease [6], dementia [7], osteoporosis [8], osteoarthritis [8], cancer [9], type 2 diabetes [10], idiopathic pulmonary fibrosis [11].

Three categories of aging hallmarks have been identified: (1) primary, or the sources of age-related harm; (2) adversarial, or the responses to the damage; and (3) cohesive, or the results pertaining to the responses and factors of the aging phenotype. Senescence, a biological process that limits the proliferation of aging or damaged cells [12], is categorized under the antagonistic category. Senescence, while serving physiological functions in proper development and maintaining tissue homeostasis, acts like a response to external stimuli elicited from age-associated damages such as genome instability and telomere loss, these are fundamental traits of senescence. There exists a profound connection between senescence and the contrasting traits associated with aging. For instance, aging cells have diminished mitophagy, resulting in “aged,” impaired mitochondrial mechanism and may contribute to metabolic disorders associated with aging [13]. Senescence additionally impacts the comprehensive characteristics of aging. Somatic multipotent stem cells promote tissue equilibrium; like, HSCs (hematopoietic stem cells) regenerate the circulatory system. Stem cell depletion transpires along aging, resulting in diminished stem cell efficacy and their regenerative potential, which contributes to tissue degeneration [14].

Following successive serial passages in culture, mesenchymal stem cells cease to proliferate post specific number of passages. Upper limit for population doublings (PDs) attainable by MSCs is projected to range between 30 to 40 depending on the tissue of origin [13]. There is no explicit information regarding the passage numbers for the 15 MSC products that have been authorized for clinical applications yet. Nevertheless, considering that the differentiation capacity of MSCs diminish following prolonged passages, it is advisable to utilize early-passage cultures for large-scale production of MSCs in bioreactors before the MSCs reach their Hayflick limit [14].

Cellular senescence and stemness have always been seen as biological opposites. Senescent cells irreversibly cease division due to cellular stress, hence inhibiting the proliferation of compromised cells, whereas stem cells facilitate tissue regeneration by perpetually renewing and producing specialized cells. Nonetheless, increasing data indicates that the interplay between these two biological processes is far more intricate and contingent upon tissue type, physiological circumstances, and disease setting. The antagonistic interaction between stemness as well as senescence: Numerous biological environments are recognized from where the initiation of senescence appears to inhibit stem cell functionality, indicating a competitive relationship between these two phenomena. An illustration of this is MSCs (mesenchymal stem cells), due to multipotent characteristics, are seen as potential agents for tissue restoration as well as several clinical applications, including repair of cartilage, immunological action regulation. Numerous advantageous traits of MSCs are linked to their capacity to develop into enhanced density single layers referred to MSC sheets, that can, swiftly exhibit signs of senescence under prolonged high-density circumstances. The mechanistic Target Of Rapamycin (mTOR) is the principal regulator for cellular functions including development, protein production, and lipid synthesis. Research demonstrated rapamycin, a prominent mTOR inactivator, enhanced autophagy of PMSCs (placental-derived MSCs), prevented the onset of senescence, which was associated with a reduction in cell size. Consequently, rapamycin was suggested as a viable approach to inhibit senescence as well as enhance use of MSC monolayers [15].

The synergy between stemness and senescence: While opposite effects among the two is noted for various tissues in normal biological states, including adipose tissue, bone marrow, pancreatic islets, and muscle, effective coordination is predominantly observed in realm of tumorigenesis. Stemness-related senescence seems to first conceal developing stem cell characteristics in a senescent cell; yet, these characteristics might eventually play a role in the “escape from senescence” phenomena.

### 1.1 Multiple ways to assess senescence

Among the various forms of cellular senescence identified, replicative senescence denotes the progressive decline in the ability to proliferate following a finite number of cellular divisions. Repeated cellular divisions culminate to shortening telomeres, activating the DNA damage signals that causes cell cycle cessation through upregulating various cell cycle inactivators, such as p16INK4a as well as p21CIP1 [20].

#### 1.1.1 Morphological alterations

Cells have several morphological anomalies during senescence, characterized by an expanded and flattened shape, along with augmented mass of multiple organelles, including lysosomes, mitochondria, and the nucleus [21]. The dimensions of cellular structures, weight of membrane organelles gradually augment throughout cellular senescence [22]. Increased cell dimensions subsequently lead to cell cycle cessation linked to senescence [23]. The dimensions and granularity of the cells can be measured both by microscopy and flow cytometry [23]. Aging cells have compromised nuclear stability owing to reduced expression of lamin B1, which acts as a hallmark for cellular senescence. Expression of Lamin B1 is usually quantified using western blot and qRT-PCR techniques. In senescence associated β-galactosidase activity, β-D-galactosidase in the lysosome hydrolyzes β-associated terminal galactosyl moieties [24], exhibiting peak activity within the pH range of 4.0 to 4.5. β-galactosidase is present alongside elevated lysosomal content in aging cells [24] and may be detected at a pH of 6.0. Histochemical as well as cytochemical activity of senescence-associated β-galactosidase (SA-β-gal) is assessed utilizing chromogenic β-gal substrate (5-bromo-4-chloro-3-indolyl-β-D-galactoside (X-gal)) along with a solution of pH 6.0 followed by fixing of cells. Quantified by measuring fluorescence post treatment by fluorescein di-β-D-galactopyranoside (FDG) or 5-dodecanoylaminofluorescein di-β-D-galactopyranoside (C_12_FDG); β-gal substrates release fluorescence (excitation wavelength: 488 nm, emission wavelength: 530 nm) when cleaved by the enzyme, which can be detected by conventional flow cytometry. [25]. To understand and observe changes associated with basal level lysosomal dysfunction that naturally characterize senescing cells various agents are used. To measure cellular senescence, L-leucyl-L-leucine methyl ester (LLOME), a specialized lysosomotropic agent, is used for initiating lysosomal membrane permeabilization (LMP), subsequent organelle damage. Late-stage autophagy inactivators, like chloroquine and bafilomycin were used in modulating lysosomal stress [42].

#### 1.1.2 Cell Cycle Arrest

The best recognized feature in senescence, cell cycle arrest. Cessation of the cycle is marked through increased expression in cyclin-dependent kinase (CDK) inactivators like p16INK4a and p21CIP1. The engagement of principal reactions, p16INK4a/RB and p53/p21CIP1, inhibits cellular growth, hence facilitating senescence. p16INK4a produced from CDKN2A gene, while p21CIP1 is synthesized from the CDKN1A gene. During senescence, p16INK4a levels rise due to different cellular stressors, while p21CIP1, elevated by p53 activation through phosphorylation in response to damage in the DNA [26].

#### 1.1.3 Senescence-Associated Secretory Phenotype SASPs

Senescent cells release interleukins, cytokines, and growth promotors, together called senescence-associated secretory phenotype (SASP). These components affect adjacent cells by modulating several cell-surface receptors and signaling cascades [32]. It has been observed that approximately 30% to 70% of senescent cells exhibit diverse SASPs, marked through secretion of cytokines which are pro-inflammatory, pro-apoptotic and immunomodulatory like IL-1, IL-6, IL-8, IL-13, IL-15, interferon-β (IFN-β), transforming growth factor β (TGF-β1) to name a few, chemokines like Eotaxin 1–3, MCP-1/CCL2, MCP-2/CCL8, tissue-degrading proteases like matrix metalloproteinases (e.g., MMP1, MMP3, MMP9), ligands (e.g., Fas ligand), growth factors [e.g., insulin-like growth factor 1 (IGF-1), IGFBP4, IGFBP7, hepatocyte growth factor (HGF)], among others [33]. Studies have progressed for detection techniques as well as genetic engineering has allowed for the identification of essential elements within the Wnt, IGF1, transforming growth factor-β (TGF-β), plasmin, signaling of interleukin (IL) pathways, thereby aiding in the control of factor release. Furthermore, SASP is pivotal in the pathogenic functions of senescent cells [21]. Quantitative reverse transcription polymerase chain reaction (qRT-PCR) and antibody arrays are frequently employed for the identification of SASP expression.

### Other advanced methods

#### 1.1.4 Microscale Magnetic Resonance Relaxometry (μMRR)

The research established a swift, non-invasive technique for identifying cellular senescence in MSCs through microscale magnetic resonance relaxometry (µMRR). The investigators quantified the T₂ relaxation time linked to intracellular iron buildup and contrasted it with recognized indicators of aging. They assessed MSC cultures via various passages, donors, cell sizes isolated via spiral microfluidics, and conditions of drug-induced senescence. The findings indicated a robust association between T₂ relaxation periods and indicators of senescence, illustrating the technique’s capacity to detect senescent cells throughout diverse populations. This method has the potential to enhance quality assurance in cell therapy production through the facilitation of live-cell evaluation [34].

#### 1.1.5 Prelamin A accumulation

Bellotti and colleagues examined if the increase of prelamin A may act as a dependable indicator of senescence MSCs. Investigators analyzed three principal human MSC lines at various phases of their in-vitro longevity by immunostaining. They evaluated prelamin A in conjunction with recognized senescence indicators, like SA-β-gal, p21, as well as Ki67. Cells in the late passage exhibited elevated prelamin A accumulation, indicative of senescence. Cells positive for prelamin A also exhibited p21 expression and were devoid of Ki67, signifying cell-cycle arrest. The results endorse the use of prelamin A detection as an effective method for evaluating MSC quality prior to clinical usage [35].

### 1.2 Image Analysis in aid of morphometric properties of cells

#### 1.2.1 Senescent cells have measurable morphological changes

One of the initial and most beneficial justifications is that senescent cells exhibit distinctive morphological alterations, such as cellular hypertrophy, flattening, augmented nuclear size, and modified nuclear/cytoplasmic arrangement. A traditional imaging-cytometry investigation reported that the flattening of senescent cells can be measured via nuclear morphology and fluorescence intensity. The authors suggested that the nuclear area and DNA-associated fluorescence might offer a quantitative assessment of the “depth” of senescence, rather than merely categorizing it as senescent or non-senescent [36]. Image analysis transforms a qualitative observation of senescent cells appear larger and more flattened into quantifiable metrics including: cell area, nuclear area, nuclear/cytoplasmic ratio, nuclear intensity, cell morphology, and cell density. This renders morphology a measurable aspect of a senescence phenotype.

#### 1.2.2. Senescence cannot be accurately determined using one specific marker

This is likely the most compelling rationale for image-centric evaluation. At present, there is no universally recognized indicator for cellular senescence. Typical indicators including SA-β-gal, p16, p21, diminished EdU incorporation, and SASP components each reflect various facets of the phenotypic and may also manifest in alternative biological conditions. The current FAST research holds significant relevance in this context. Neri et al. devised an automated image-analysis technique that concurrently measured: SA-β-gal expression, uptake of EdU /proliferative cessation, along with nuclear enlargement at the individual cell level. The writers highlight that integrating various data yields a more comprehensive evaluation of senescence compared to depending on a solitary marker [37]. This provides an individual with a solid justification that image analysis facilitates the concurrent measurement of various independent characteristics of senescence within a single cell. This is significantly more enlightening than assessing, for instance, the overall β-galactosidase activity within a population.

#### 1.2.3 Image analysis reveals cellular diversity

Senescent populations exhibit heterogeneity. Two cells subjected to identical senescence-inducing stimuli may have markedly varied levels of SA-β-gal, p21, p16, nuclear enlargement, or SASP expression. A review from 2022/2023 emphasizes the significance of examining senescence at the single-cell level due to considerable variability of genetic as well as functional traits across senescent cells [38]. Recently, Seshadri et al. employed single-cell fluorescence imaging to quantify SA-β-gal, p21, IL-6, nuclear area, and cell area. They discovered considerable variability in nearly all of these characteristics. Notably, specific subpopulations within the nuclear region were linked to variations in IL-6 expression [39]. This represents a significant benefit associated with image analysis, a standard plate reader or western blot may indicate that the culture exhibits elevated p21 levels. But through image analysis one can understand the percentage of cells that exhibit elevated p21 levels, however this group comprises multiple phenotypically diverse subpopulations. That differentiation may hold biological significance.

#### 1.2.4 Image analysis facilitates the quantification of senescence at the single cell level

The intricately linked heterogeneity yet warrants its own distinction. High-content imaging enables each cell to serve as a distinct data point. Consequently, one can compute percentage of senescent cells, intensity of specific senescence indicators, distribution of cellular/nuclear dimensions, populations that are positive vs negative for markers, and the correlations among various markers inside single cells. The research conducted by Debacq-Chainiaux and associates revealed that imaging cytometry can objectively assess morphological and molecular characteristics linked with senescence in single cells [36]. Similarly, Wang et al. formulated a quantitative digital image-analysis method for SA-β-gal staining and demonstrated that image analysis could identify relatively subtle variations in staining intensity linked to distinct senescence mechanisms [40].

#### 1.2.5 Image analysis can differentiate the extent of senescence instead of only indicating presence or absence of the biological stage

Senescence is frequently regarded as a dichotomy: senescent vs non-senescent. However, from a biological perspective, the situation is more intricate. Imaging facilitates the quantification of continuous variables like nuclear region, cellular region, SA-β-gal expression level, p21 expression level, EdU expression level. The literature on imaging cytometry has explicitly suggested that morphological metrics serve as indications of senescence depth [36].

### 1.3 Open-Source software tools used in image analysis for Stem Cells Research

In the past ten years, considerable progress has been made in realm of bioimage analysis, nascent computational discipline that collaborates closely with life sciences for the quantitative evaluation of scientific imaging. In numerous instances, significant resources have been devoted to transforming these novel advancements into functional software applications, enabling users to consistently employ state-of-the-art techniques for their analytical challenges through tools like ilastik, CellProfiler, Fiji/ImageJ2 along with its various contemporary plugins that enhance the BigDataViewer framework, among others. Such software applications have now integrated into an essential framework for scientific endeavors.

**Table 1:**
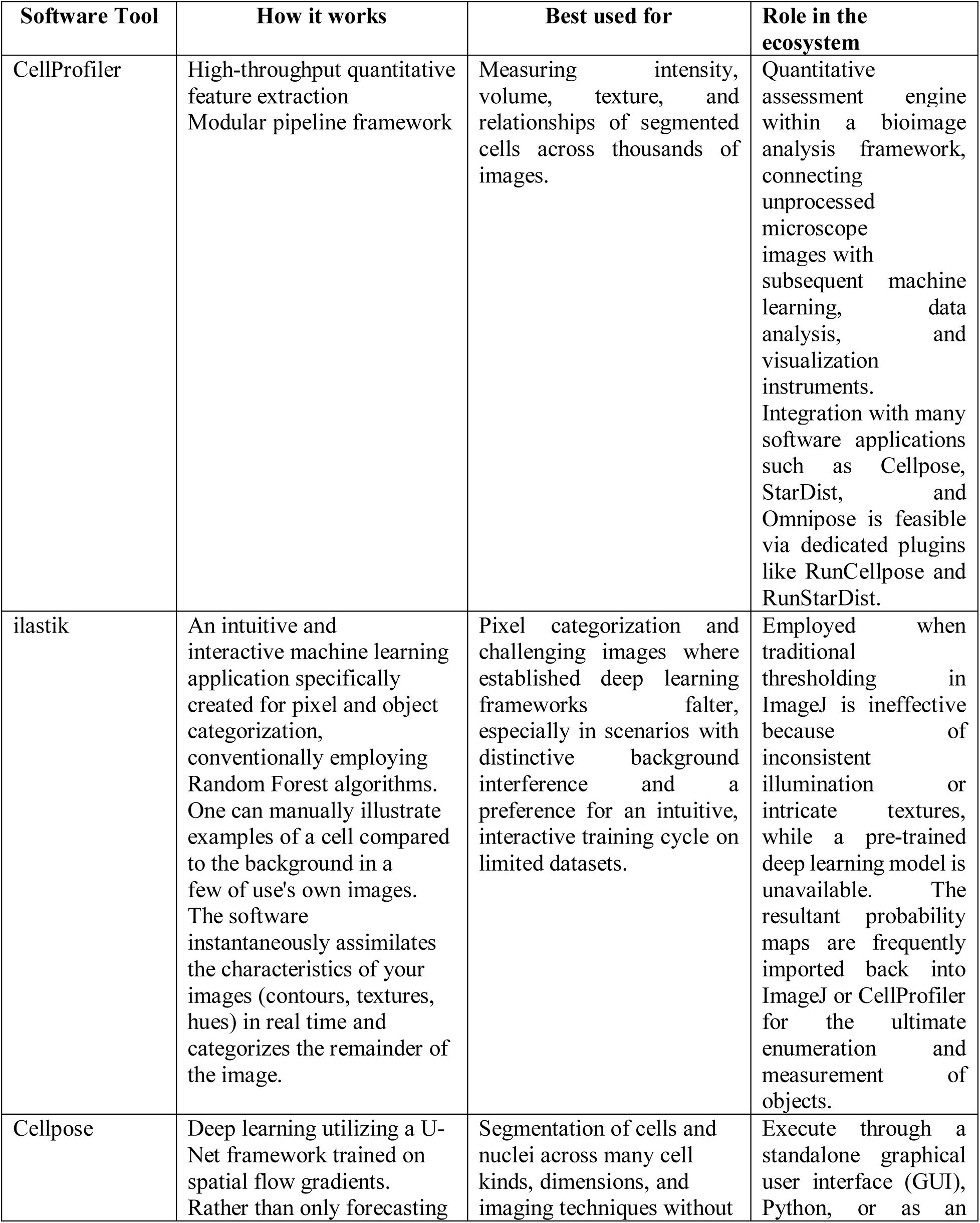
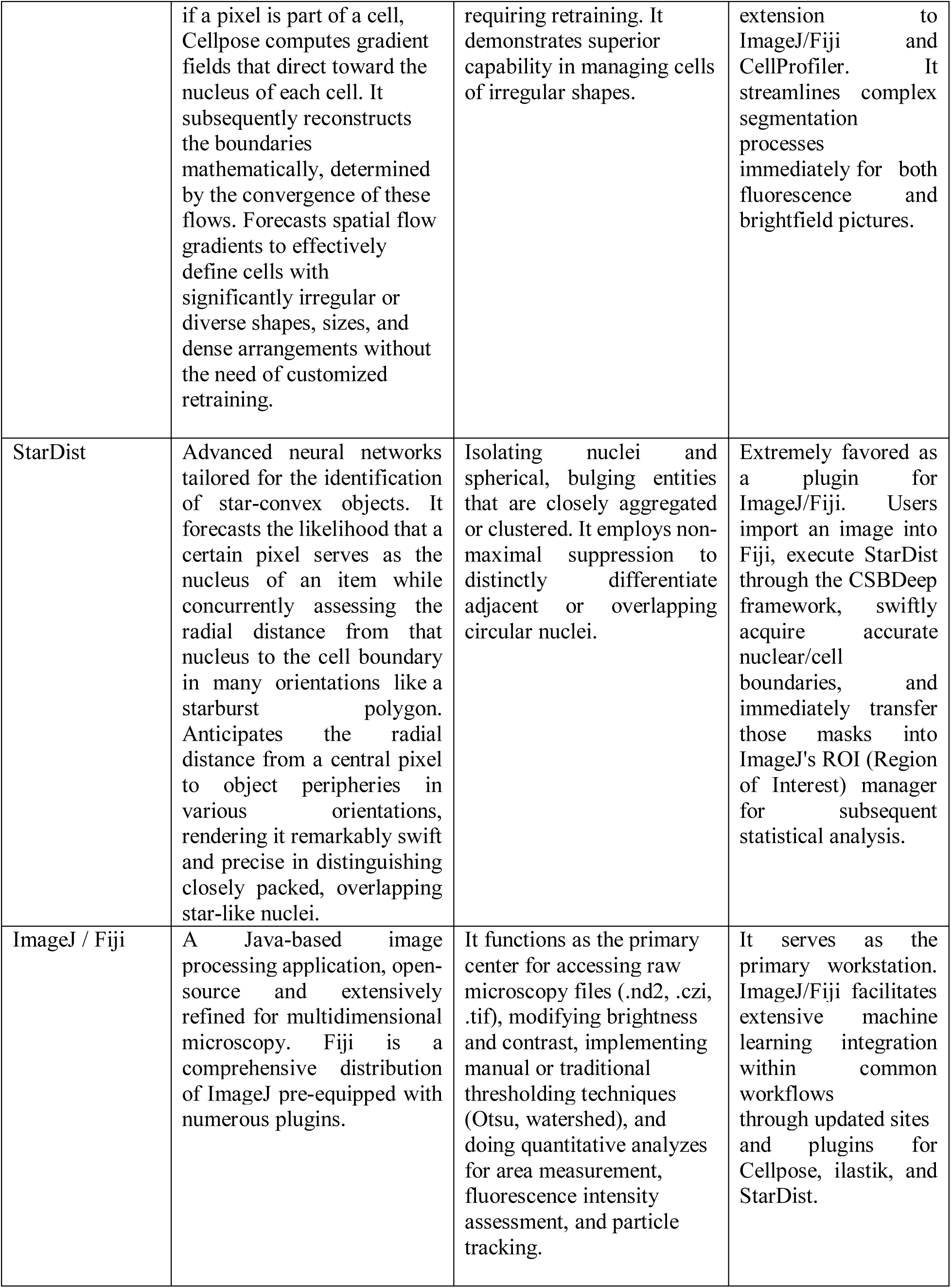
Software tools used to analyze image data in biological samples:

#### 1.3.1 MSCProfiler a novel tool used to study heterogeneity in MSCs

MSCProfiler is an automated approach for single-cell image analysis created with CellProfiler software to examine the morphological and immunophenotypic diversity of mesenchymal stem cells (MSCs). Gupta et al. in 2023 [41] described an analysis of brightfield and fluorescent images acquired with imaging flow cytometry. The pipeline retrieves quantitative characteristics, encompassing cell area, morphology, eccentricity, compactness, texture, entropy, and fluorescence intensity of surface markers like CD44 and CD73.

Its primary advancement lies in the amalgamation of automated image quality assessment, cellular segmentation, morphological and textural feature extraction, along with biomarker intensity quantification into a cohesive process. MSCProfiler facilitates the relatively unbiased and high-throughput analysis of individual cells by eliminating subpar or irrelevant image files and minimizing manual input. The research proved its effectiveness in recognizing diverse MSC populations from various tissue origins and culture stages, enhancing traditional flow cytometry. MSCProfiler is especially advantageous for scrutinizing extensive imaging flow cytometry datasets while exploring MSC heterogeneity, contrasting cellular morphology across passages or tissue sources, and correlating morphological patterns with biomarker expression. Exported data can facilitate statistical analyses and subsequent classification through machine learning. To achieve best outcomes, users must set suitable image-quality benchmarks, verify segmentation and fluorescence assessments, and manage datasets in feasible increments. The primary constraints are the computational requirements and the necessity for supplementary analytical instruments when categorizing cells based on multiple characteristics.

Moving forward from this work established with MSC profiler for MSC heterogeneity we have built a pipeline with modifications to the MSC profiler specific to assess MSC senescence properties through bright field and fluorescent input images. The steps followed in the modified pipeline is given in detail in the next section.

## 2. Materials and Methods

### 2.1 Cell Culture and maintenance

Human Dental Mesenchymal Stem Cells were maintained and cultured at specific passages of P6, P9 and P12 for this study. The cells were seeded in 6 wells plates, grown in Dulbecco’s modified Eagle medium (Gibco) media supplemented with Sodium pyruvate (Gibco), Sodium bicarbonate (Himedia) and 10% fetal bovine serum (Gibco), 1X Pen-Strep (Gibco), allowed to grow in 37°C (humidity) to reach a 75–80% confluency and media changed every 24 or 48 hours.

### 2.2 Senescence associated β-Galactosidase activity through Chloroquine diphosphate

Once the cells reached a confluency of 75-80%, the monolayer was washed with Dulbecco’s Phosphate Buffer Saline (Himedia) and 1ml of media was added to each well. To the test well 45μM chloroquine diphosphate (Himedia) was supplemented and placed at 37°C for 90 mins following which to the same well 33μM C_12_FDG (Invitrogen) was supplemented and placed for another hour at 37°C. All other required controls along with single colours used as a technical requirement for the imaging flow cytometer were kept during the experimentation

### 2.3 Preparing single cell suspension

The monolayer was washed with Dulbecco’s Phosphate Buffer Saline and 0.05% Trypsin-EDTA (Gibco) was added and incubated at 37°C was the cells to detach. Trypsin-EDTA was neutralized with Dulbecco’s Phosphate Buffer Saline, cells were collected in FACS tubes and centrifuges at 200g for 2 mins at room temperature to obtain a pellet. The supernatant was discarded and pellet was resuspended in staining buffer (2% FBS in Dulbecco’s Phosphate Buffer Saline) and 0.01μM of DRAQ-7 (Invitrogen) was added. Immediately acquired on Amnis Image Stream MK II Imaging Flow Cytometer.

### 2.4 Acquisition on the Amnis Image Stream MK II Imaging Flow Cytometer

Technical requirements are single colour green and red tubes which will be used for compensation post-acquisition.

Single colour green: the cells are treated with 45μM chloroquine diphosphate and incubated at 37°C for 90 mins followed by 33μM C_12_FDG was added and incubated for one hour at 37°C. Monolayer was washed with Dulbecco’s Phosphate Buffer Saline and 0.05% Trypsin-EDTA was added and incubated at 37°C was the cells to detach. Trypsin-EDTA was neutralized with Dulbecco’s Phosphate Buffer Saline, cells were collected in FACS tubes and centrifuges at 200g for 2 mins at room temperature to obtain a pellet. The supernatant was discarded and pellet was resuspended in staining buffer.

Single colour red: Cells were washed with Dulbecco’s Phosphate Buffer Saline and 0.05% Trypsin-EDTA was added and incubated at 37°C was the cells to detach. Trypsin-EDTA was neutralized with Dulbecco’s Phosphate Buffer Saline, cells were collected in FACS tubes and centrifuges at 200g for 2 mins at room temperature to obtain a pellet. The supernatant was discarded and pellet was resuspended with 30% hydrogen peroxide and incubated at room temperature for 30 mins in dark. 2ml of Phosphate Buffer Saline was added and centrifuged at 200g for 5 minutes to obtain a pellet. The pellet was resuspended in staining buffer and 0.01μM of DRAQ-7 was added.

#### 2.4.1 Building compensation matrix in Ideas software

Conventional flow cytometry detects voltage gain of individual photomultiplier tubes (PMTs) upon each fluorescent signal of the antibody fluorochrome conjugate. This is not the case for imaging flow cytometry (IFC) as the detector is a charged coupled detector (CCD) chip which has higher quantum efficiency than PMTs and can detect dimmer signal required for detecting maximum information in an image. It provides an output as an image and requires pixel level compensation. To perform pixel level compensation “noBF” images were acquired as single colour control which were used to build the compensation matrix. The resulting compensation matrix was saved in .ctm format.

#### 2.4.2 Data Analysis and Gating strategy

Post compensation two more files are formed along with the raw image file, one is “. cif” and the other is “.daf” file. The .daf or the data analysis file is what is used for analyzing the acquired data. Each dot in the imaging flow cytometry plot represents image of each cell. We sequentially gated the live senescent images (figure 2) by assigning regions in each plot. Starting from the first histogram that gates out focused cell images using gradient root mean square value. As sharp change in pixel represents boundary of highly focused images hence, higher the gradient RMS value higher is the focus. From the focused images we select the single cells using area vs aspect ratio plot assuming single cells will have lower area and aspect ratio near a perfect circle i.e., 1. From the single cells we gate out live or dead cells using live dead marker such as Draq 7 and them from the live cells we gated the higher β-galactosidase as region as High and for the lower expression below 10^5^ intensity as low region correlating with the corresponding image. In figure 3 we show the images acquired in each channel and where channel 1 is brightfield (BF) image, channel 2 is the image for β-galactosidase expression, channel 5 is for Live Dead marker (Draq 7/ To - Pro) and channel 6 shows the side scatter/ dark field image.

### 2.5 Modified MSCProfiler workflow

The steps of the pipeline and their result is discussed in figure 4. In this section we will walk through each of those steps in detail.

Quality control – As a quality control measure any image above the intensity of 4000 were allowed to pass to the downstream steps of the pipeline using the “FlagImage” module. “MeasureIntensity” module was used that measure the intensity using the bright field (BF) channel (Ch1). This allows to find out whether a cell is present in an image or not.

Preprocessing – The unflagged images by the “FlagImage” module was used for further analysis. The BF channel image was preprocessed to make it easier for the pipeline to annotate a single cell as foreground. Steps such as background subtraction, smoothening, rescaling intensity, edge detection have been taken for the preprocessing process. Background subtraction was done using “ImageMath” module. Then the image was inverted using “Image Math” module rescaled using “RescaleIntensity” module which rescaled every pixel, the minimum intensity was 0 and the maximum intensity was 1. Post rescaling the “EnhanceEdges” module was used to detect the edges.

Segmentation of the cell boundary – This process was done to identify the cell from each image. The processed image was smoothened that reduced the contrast within the edged image of the cell. This made segmenting the entire cell as foreground feasible. Segmentation of the entire cell was done using the “Threshold” module by Otsu method based on pixel value variation between foreground and background. “IdentifyPrimaryObject” module was used to identify the cell boundary. The cell boundary was adjusted using the “ExpandorShrinkObject” module enhancing the boundary detection accuracy. Any identified object whose boundary was touching the boundary of the image or having area below 2500 pixel were flagged and not allowed for downstream processes.

Segmentation of Channel 2 fluorescent image - The input images were rescaled using “Rescale Intensity” to brighten the dimmer pixel. Smoothening was done using the “Smooth” module that reduced the pixel variation in the fluorescent image. “IdentifyPrimaryObject” was used to segment the fluorescent cell boundary in channel 2.

Segmentation puncta in channel 2 – To make the high intensity pixel identifiable both by human eye and the algorithm “EnhanceandSuppressFeature” was used that helps enhance or suppress certain identifiable features within the image like speckles, neurites etc. Here we enhanced speckles and segmented the brighter spots using minimum cross entropy thresholding technique in “IdentifyPrimaryObject” module. We identified the punctate morphology for 1210 images automatically using this module across three biological repeats for early, mid and late passage.

Relate each punctum to each cell – To relate identified number of puncta in an image to its bright field morphology we used “RelateObject” module. This assigns relationship of the bright field features of a cell to the number of puncta identified in it. Thus, morphological and geometrical changes can be correlated to the number of puncta henceforth replicative senescence.

Measurements of Intensity and Geometrical features – 22 different types of image intensity values and 54 morphological and geometrical features were measured for multiple images using “MeasureIntensity” and “MeasureObjectSizeShape” module for BF, fluorescent channel 2 of each cell and for each punctum.

Visualization of the segmentation – “OverlayObjects” module was used to visualize BF segmentation and puncta segmentation. Each segmented object was assigned different colors to visualize them as separate objects. “OverlayOutline” was used to visualize the outer boundary of each cell and punctum allowing the user to visualize whether the segmentation is correct or not. Save visualization and quantification result – Quantification of each image feature were saved in the form of .csv format using “ExportToSpreadsheet” module. Visualization results were saved using “SaveImages” module.

### 2.6 Western blot to validate the expression levels of p53 across different passages of hDPCS and its correlation with senescence

Samples were prepared across passage of hDPSCs with condition of untreated, chloroquine diphosphate + C_12_FDG treated, chloroquine diphosphate treated and C_12_FDG treated being constant. hDPSCs were seeded in 6 well plates and once they reached 75-80% confluency there were treated with different conditions as mentioned above.

#### 2.6.1 Protein Extraction and Quantification

1. Once the specific treatments and incubations were done 100μl of lysis buffer was added (RIPA buffer+ PIC)
2. The monolayer was scraped with the help of a cell scraper and pipette mixed for mechanical lysis.
3. The lysates were collected and incubated on ice for 20 mins, centrifuged at 14,800rpm for 20 mins at 4°C
4. The supernatant was carefully collected into fresh tubes and stored in −20°C
5. Protein quantification was done by BCA assay-BCA standards 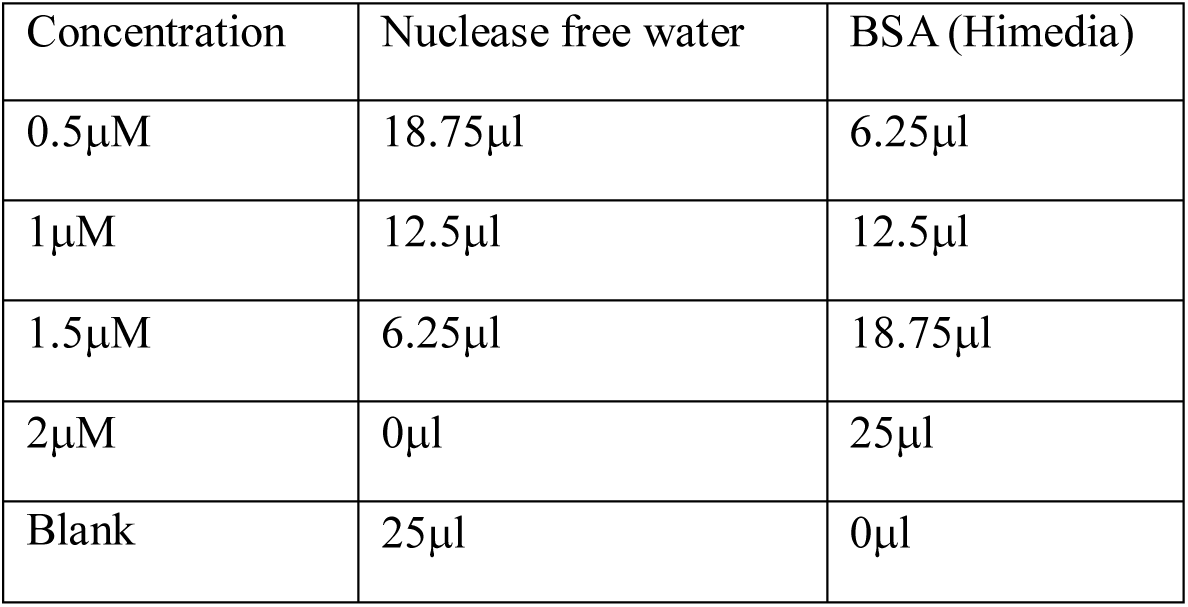 Test samples: 2.5μl of test protein lysate+ 22.5μl nuclease free water was added into a 96 well plate and to all the standards and test wells 200μl of BCA solution (BCA and CuSO4 49:1 ratio, E bioscience) was added and the plate was incubated at 37°C for 30 minutes.
6. Absorbance was recorded in a spectrophotometer at 562nm.

#### 2.6.2 Sample preparation and gel electrophoresis

1. After calculating the R^2^ values for each sample from the absorbance values, specific volumes of each sample were taken such that 10μg of protein was loaded into each well.
2. Specific volume of protein lysates were taken into fresh tubes and equal volume of laemmli buffer was added into the tubes and boiled at 95°C for 10-15 mins
3. The gel was set with 10% resolving and 4% stacking and the samples were loaded along with the ladder.
4. Once the dye front reached the bottom of the tank the gel was prepared for transfer

Table: Composition of Resolving and Stacking gel:

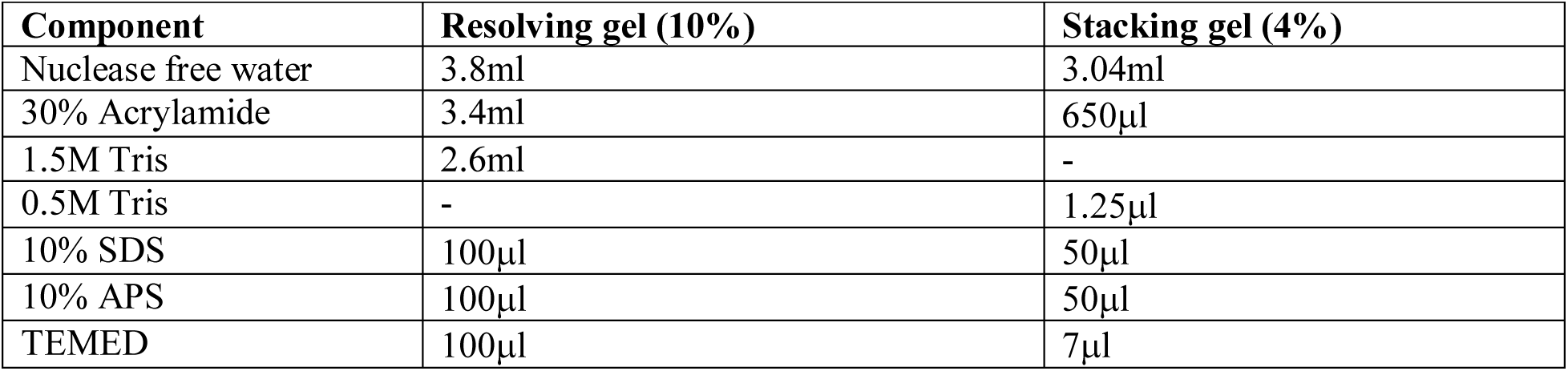

| Component | Resolving gel (10%) | Stacking gel (4%) |
| --- | --- | --- |
| Nuclease free water | 3.8ml | 3.04ml |
| 30% Acrylamide | 3.4ml | 650µl |
| 1.5M Tris | 2.6ml | - |
| 0.5M Tris | - | 1.25µl |
| 10% SDS | 100µl | 50µl |
| 10% APS | 100µl | 50µl |
| TEMED | 100µl | 7µl |

Table: Composition of running buffer (5X, 250ml):

| Components | Quantity |
| --- | --- |
| Tris | 3.75g |
| Glycine | 18.55g |
| SDS | 1.25g |

#### 2.6.3 Protein Transfer

1. Proteins were transferred from the gel to a PVDF membrane using a tank for wet transfer system (Biorad).
2. The PVDF membrane was activated in 100% methanol for one minute, followed by 1-2 rinses in transfer buffer.
3. Membrane was equilibrated in 1X transfer buffer along with the filter papers.
4. The transfer sandwich was assembled from the cathode (negative) to the anode (positive) in the pattern of fiber pad, filter paper, gel, membrane, filter paper, and fiber pad
5. The air bubbles were gently removed by a roller.
6. Electrophoretic transfer was performed at 100 V for 1 hour at 4°C.

Table: Composition of transfer buffer (4X, 125ml)

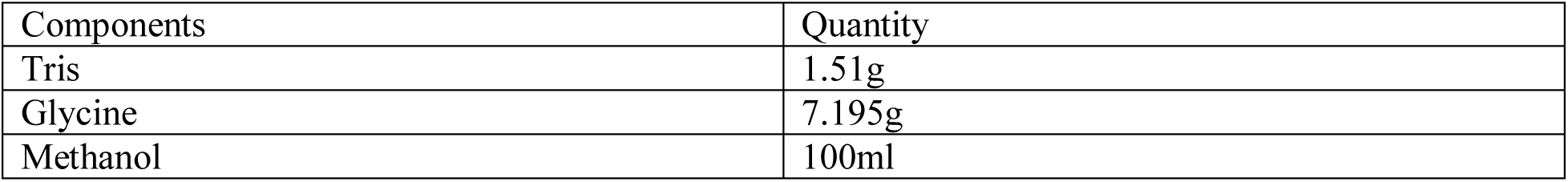

#### 2.6.4 Blocking and Antibodies

1. Once the transfer is completed, blocking solution (5% BSA in TBST) was added to the membrane and placed on a rocker at room temperature for 1 hour
2. Membrane was washed with TBST and placed on the rocker for 3 mins
3. Primary antibody (TP53, Cusabio 1:1000) diluted in blocking solution was added to the blot and placed on the rocker at 4°C over night
4. Blot was washed 2-3 times with TBST on the rocker at room temperature and HRP conjugated secondary antibody (anti-rabbit, CST) was added to the membrane and placed on the rocker for 1 hour

#### 2.6.5 Blot development and quantification

1. Blot was developed with ECL (Biorad) in the chemidoc system (Biorad).
2. ECL Solution: solution A+ solution B (1:1) freshly prepared before use
3. The solution is added on the blot near the expected molecular weight of the protein and developed in the system

<u>Quantification:</u> Band intensities were quantified using densitometric analysis in ImageJ software. The background intensity was subtracted individually from the integrated density of each TP53 and GAPDH band. The background-corrected TP53 intensity was then normalized to the corresponding GAPDH intensity for each experimental condition. The normalized values were subsequently used to generate a bar graph to compare TP53 protein expression across the different experimental conditions.

### 2.7 Differentiation of hDPSCs into osteogenic lineage

1. hDPSCs were seeded into 48 well plates and maintained in 10% media and allowed to grow in 37°C (humidity) till they reached 70-75% confluency
2. Upon reaching the desired confluency the monolayer was washed with dPBS solution
3. The controls were maintained in 2% DMEM media, and the cells to be differentiated were supplemented with StemPro Osteogenesis Differentiation Kit (Thermo Fisher Scientific, cat. no. A1007201) where the basal and supplement media are in 9:1 ratio. Incubated at 37°C
4. Media change was given every 3 days for 21 days and observed under the microscope for calcium deposit formation

## Result and Discussion

We wanted to understand the onset of senescence in mesenchymal stem cells derived from the human dental pulp while passaging them *in vitro* (n= 3). The study started by detecting the immune-heterogeneity in stem cells from human exfoliated deciduous teeth (SHEDs) and in Wharton jelly MSCs (WJMSCs). Typically, using the MSCProfiler pipeline, multiple morphometric features (size and texture) and surface biomarkers could distinguish three classes of MSCs within the same clonal population of each of SHEDs and the WJMSCs in our earlier study. To extrapolate the features for senescence in the hDPSCs, we used an experimental strategy to measure this property in stem cells in single cell suspension. To calibrate and quantify the expression of the enzymatic reaction between the beta-galactosidase and its substrate in these adult stem cells, the hDPSCs were treated with chloroquine diphosphate to increase the lysosomal pH and enhance the production of β-galactosidase which can cleave the substrate C_12_FDG supplemented in the media. Post incubation, the cells were analyzed on an imaging flow cytometer to quantify and observe the puncta formation and compare their fluorescence intensities across three different passages (early, P6-mid, P9 and late, P12). The pipeline was modified to add on the properties of puncta morphology based on their count and fluorescence intensity profiles in the green channel. Parallelly all features including cell size, shape and eccentricity from bright field images of single cells were compared across these three passages of culture.

Based on Figure 1, the authors were able to record the population of stem cells which were focused, circular in shape, live cells (based on null expression in DRAQ7 channel) and showed expression of β-galactosidase enzyme. Dot plot A was set up on the system while acquiring cells to collect only the population of cells that are focused in the middle of the stream at the point of interrogation. The focused cells were then plotted on dot plot (B) of Area vs Aspect ratio to understand the close to circularity morphology of the cells before and after the chloroquine treatment. The aspect ratio of circular cells in the range of 0.8-1 were selected for further analyses. Live cells shown in the dot plot (C) were those which did not show expression of DRAQ7, live dead discriminator. Draq 7 intensity was observed in Channel 5. Cell populations with high intensity in ch 5 were the dead cells(R4) and cells showing low intensity were the live cells (R3). The live cells were detected on the dot plot for Intensity Channel 5 vs Intensity Channel 2 (D) for estimating the cell populations for high, medium and low beta-galactosidase expression levels. We categorized the cells based on their intensity values in Channel 2 (green) and observed three classes of expression ranges. Cells treated with chloroquine diphosphate showed increased production of β-galactosidase when supplemented with the substrate C_12_FDG. The enzyme-substrate reaction showed fluorescent signal (Ex: 490/496nm; Ex: 516/525nm) collected in Channel 2 (green) of the imaging flow cytometer. We statistically classified the cells into 3 groups for expression of the intensities.

**Figure 1:**
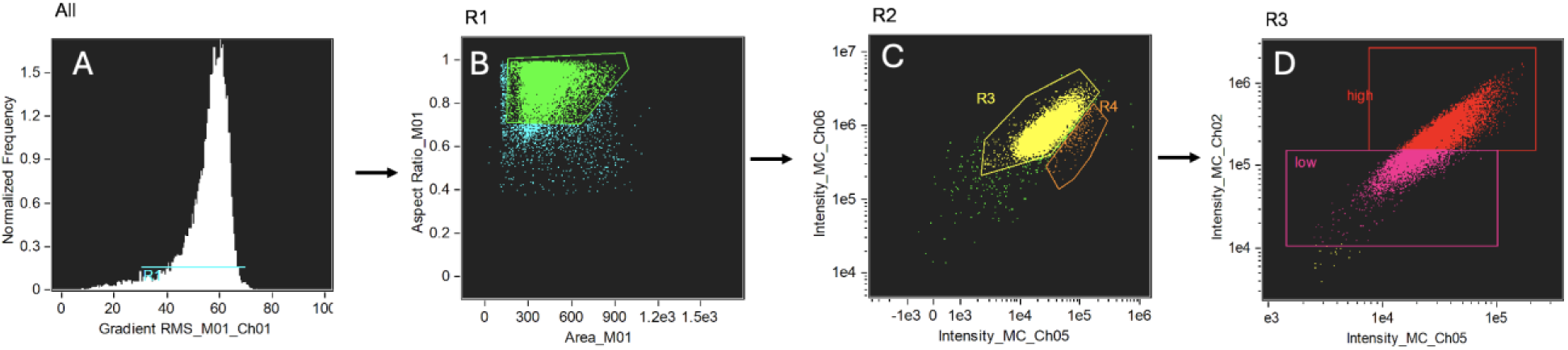
Gating strategy followed while acquiring single cells on Amnis Image Stream MK II. Plot A shows the population of cells which are focused in the middle of the flow stream. (B) Dot plot showing the cell area in terms of aspect ratio to gate circular cells. (C) Dot plot of Intensity ch 5 verses Intensity ch 6 for a live dead discrimination. (D) Dot plot of Intensity ch 5 verses Intensity ch 2 depicts cells which have high and low expression of beta-galactosidase production.

**Figure 2:**
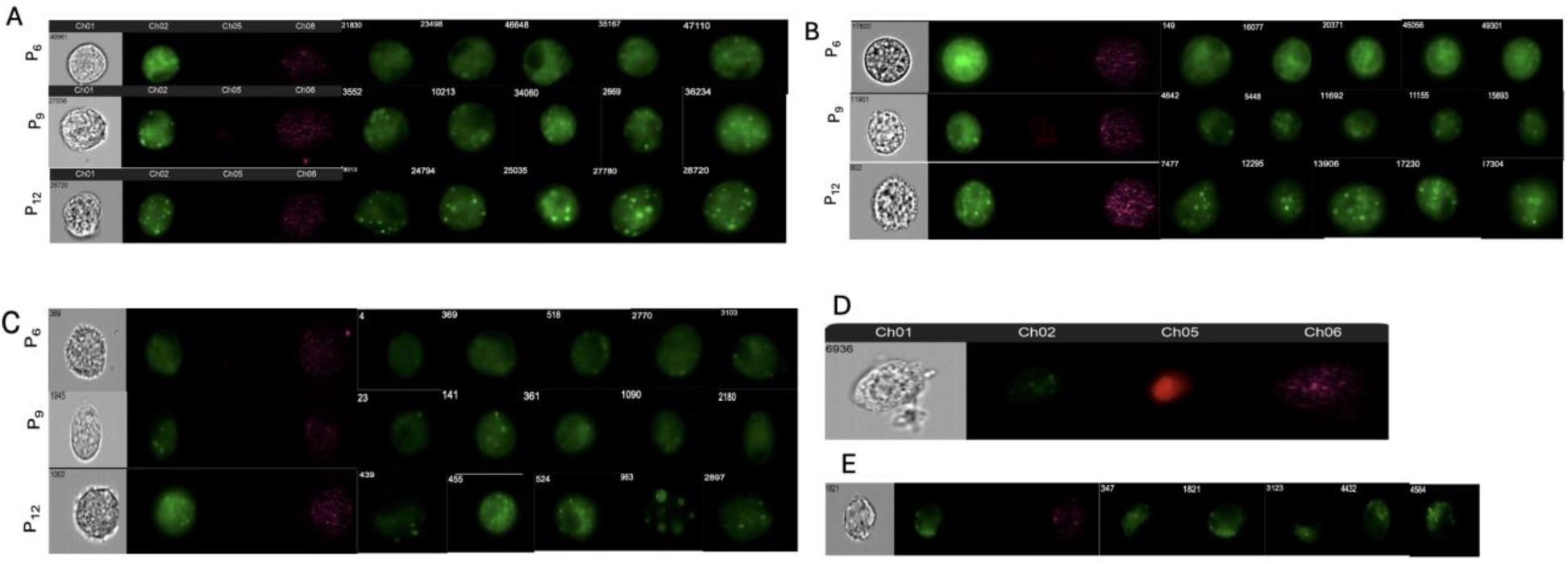
Image gallery generated from Amnis Image Stream MK II. Images taken from different biological repeats (N=3) figure (A-C) and across different passages (P6, P9 and P12). (D) Dead cell image which has taken upon the dye. (E) HeLa was the control used for this study

The above gallery of images in Figure 2 represents the image gallery of cells which were analyzed using the IDEAS software. This study required the Amnis ImageStream MK II having a single camera, 3 laser configuration (405nm, 488nm and 642nm), 6 detector channels, with multiple magnification options (20X,40X and 60X) and with the extended depth of field (EDF) feature. Since the puncta morphology was the prime focus in this study for senescence, the EDF feature proved very useful for the expression levels and in generating the images.

The study was conducted with three biological repeats. Single cell fluorescent images showed an increase in the count and intensity of punctate structures as observed in the single cell images. It was clearly visible that the puncta morphology based on the expression of β-galactosidase, was different across the 3 passages of growing cells. In the early passage (P6) the punctate morphology was majorly diffused when compared to the distinct and high intensity punctate morphology seen in the mid and late passages (P9-P12). The authors observed the same trend across all the biological repeats. Using the proprietary IDEAS software, we could not distinguish between the punctate expression and found all the 3 passages showing both fuzzy and structured expression of the enzyme. We used this software to define a puncta in a single cell as dotted foci like structures, visible in the image gallery and quantifiable from the pixel values of the fluorescence intensities. This was typically how we expected to see the fluorescent enzyme-substrate reaction from the affected lysosomes as dotted structures observed under the microscope in adherent cells.

HeLa, a cancer cell line which ideally should not show replicative senescence properties were used as control cell lines in the study.

**Figure 3:**
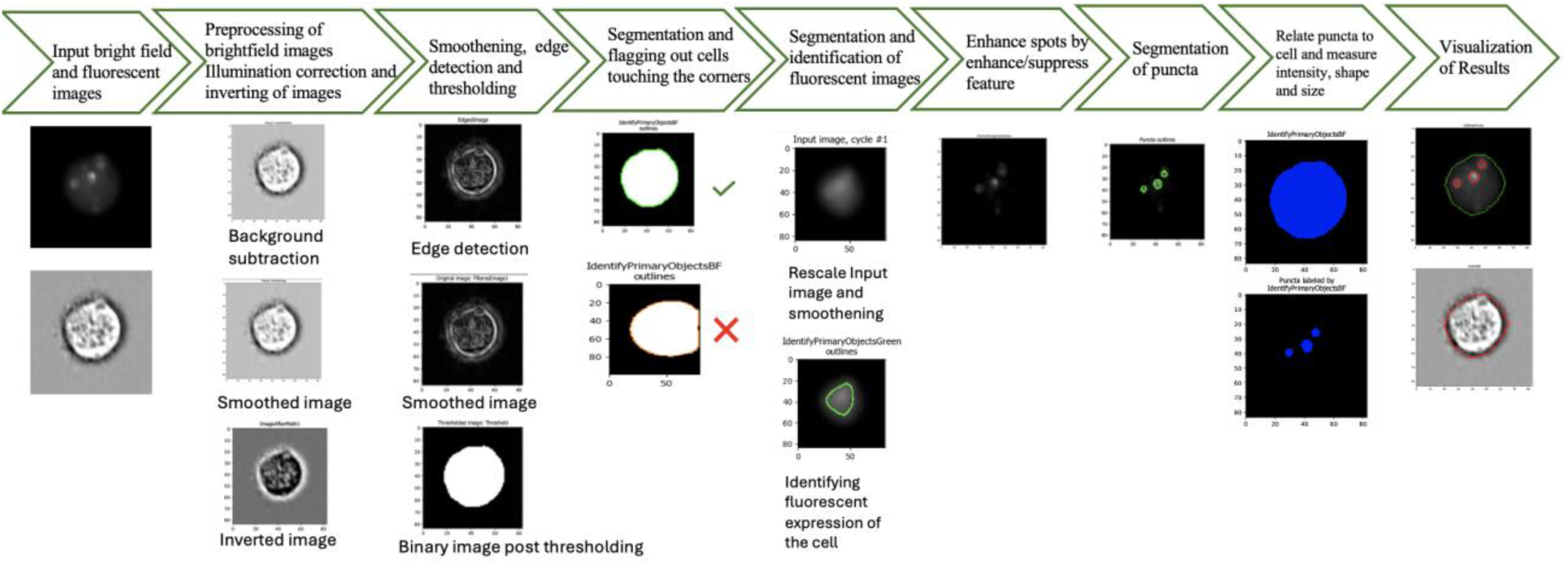
Modified MSC profiler pipeline for assessing senescence in hDPSCs. Brightfield and fluorescent images were used to analyze the number of puncta and its correlation with *in vitro* aging. The pipeline also analyzed morphological changes in senescent cells like cell size, shape and eccentricity.

Once the images were analyzed on the IDEAS software, multiple sets of brightfield and fluorescent images were uploaded onto the modified MSCProfiler pipeline to compare and quantify the punctate morphology in these hDPSCs. Figure 3 describes all the steps in the modified MSCProfiler pipeline to quantify senescence in these adult stem cells. The images undergo several preprocessing steps like illumination correction, background subtraction, smoothening and image inversion. The edges are detected and smoothened. The images are converted into binary vales after thresholding where the cell is considered as 1 and the background as 0, post thresholding the images are segmented and all the cells which are not aligned in the middle are excluded. Images showing fluorescence were processed to enhance/suppress the punctate morphology in the cells, after proper identification the puncta are segmented, its intensity, size and shape values are obtained and the results statistically interpreted. Apart from puncta identification and quantification, the pipeline also gave morphological feature descriptors like the cell size, shape, compactness, solidity and eccentricity of cells across different passages of hDPSCs.

**Figure 4:**
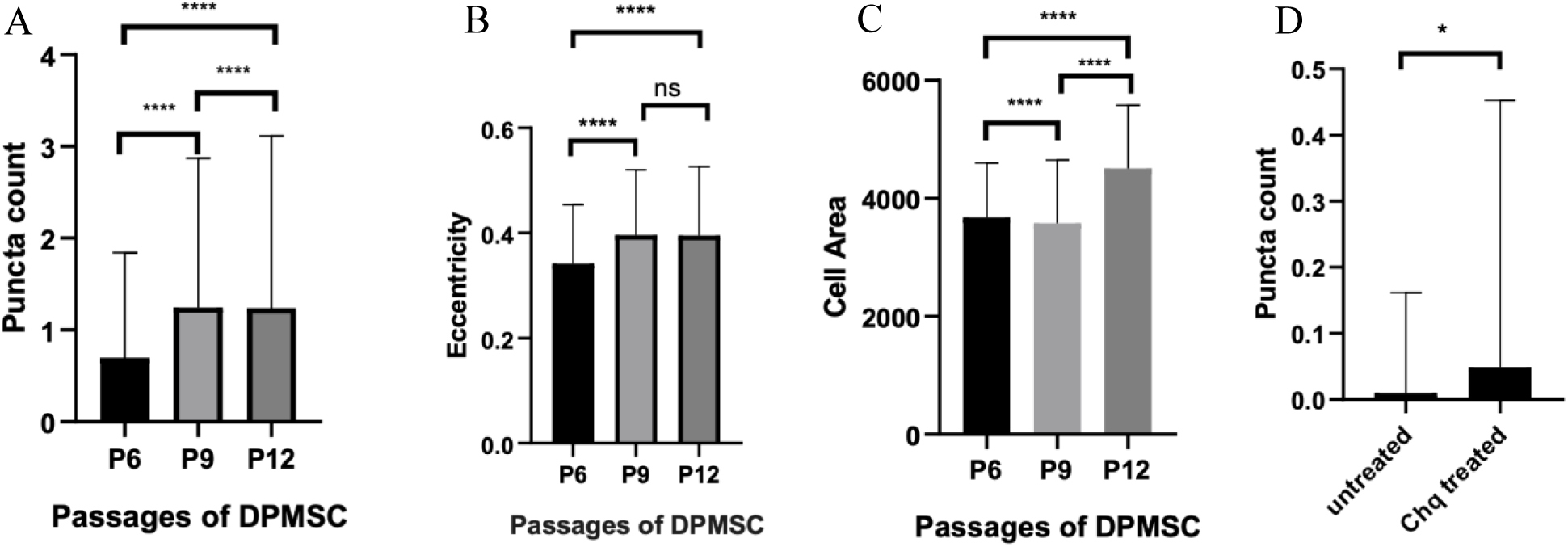
Quantified fluorescent and brightfield image features across three passages by modified MSCProfiler. All the cells used for the quantification were chloroquine treated and supplemented C_12_FDG. (A) Puncta count, (B) Eccentricity, (C) Cell area and (D) Puncta count in HeLa.

Puncta count and puncta intensity profiles were mainly used to measure senescence in cells based on the values generated by the pipeline. The aspects which were looked at were the puncta count, eccentricity and cell area across different passages in DPSCs. The authors have observed an increase in the number of puncta as the passage increases. Eccentricity for a perfect circle is zero but the data shows deviation in the eccentricity values for cells treated with chloroquine diphosphate across all the passages in DPSCs. Authors have also seen an increase in the overall cell area across passages in senescent cells. HeLa cell line on the contrary were analyzed in the same manner and there was no significant difference between the treated and non-treated population of these cells.

**Figure 5:**
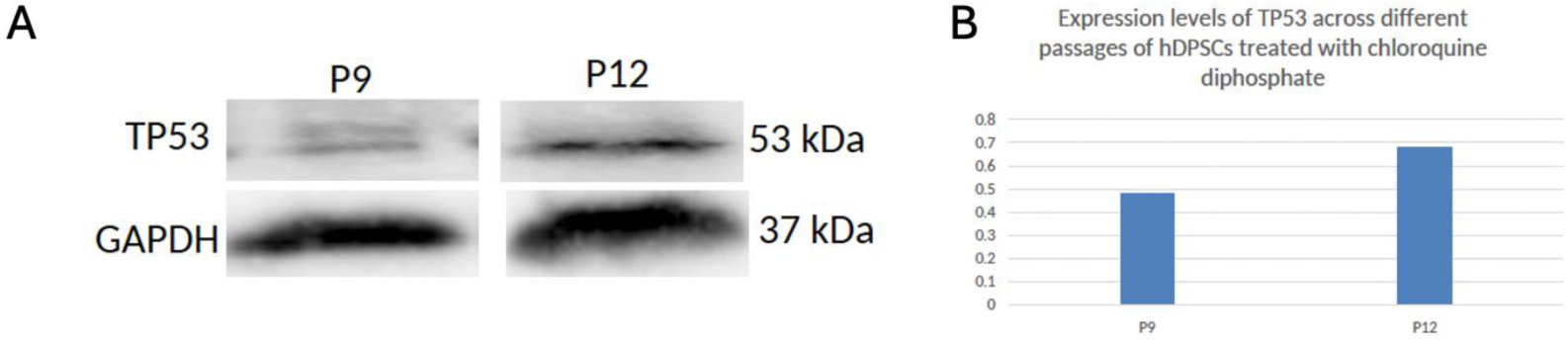
Expression levels of TP53 across passages of DPSCs. The mid and late passage cells were treated with chloroquine diphosphate and checked for varying expression levels of TP53. (A) Western blot bands. (B) Quantification of the amount of protein

To validate that the puncta morphologies were correctly correlating to the senescence properties of the hDPSCs across the 3 passages, we wanted to quantify a definitive marker of a senescing cell using the western blot assay. We optimized the expression levels of TP53 protein in the cells which were treated with chloroquine. TP53 is upregulated in senescent cells. The expression of TP53 was found to be lower in P9 when compared to P12 upon treatment with chloroquine diphosphate, suggesting an increase in senescence as the *in vitro* age increased in these MSCs. The internal control housekeeping protein used for the assessment was GAPDH.

## Discussion

The prime focus of this study was in using an automated analysis pipeline (MSCProfiler), which had already been trained to provide image quality control right up to parameter estimation and classification of mesenchymal stem cells and plug in the senescence features of the MSCs. Post segmentation, these images were classified to measure geometrical features such as shape, size, eccentricity along with intensity values of the β-galactosidase from over 50,000 single cell images obtained from imaging flow cytometry. The results showed that β-galactosidase expression could be correlated to the puncta morphology, and it was defined for the first time in hDPSCs at the single cell level. Imaging flow cytometry enabled the detection of the puncta or foci structures appropriately with the help of the EDF feature, an important hardware requirement of the technology. The sizes could be measured, and we could use the pixel intensities to classify the low, medium and high β-galactosidase expression patterns in the cells.

Use of image analyses pipelines to quantify and measure parameters of cells have been mostly used for microscopy images. Here we modified our own pipeline MSCProfiler to analyze the imaging flow cytometry data that added valuable information from the images concurrently with quantification. The hallmark of this screening pipeline was the identification of puncta morphology from single brightfield and fluorescent images of single cells. The long term plan would be to use such a pipeline to correlate the properties of senescence with lineage differentiation of the MSCs. In this case we are currently looking at the capacity of hDPCS to differentiate into the osteogenic lineage across the 3 different passages. Such a pipeline can be used in future as a CQA specific to MSCs that can predict the age of cells before their use in downstream applications such as any kind of allogenic transplantation.

